# Smooth Muscle Cell Cytoglobin is a Negative Regulator of Atherosclerotic Fibrous Cap Development

**DOI:** 10.64898/2026.06.25.734607

**Authors:** Kurrim Gilliard, Le Gia Cat Pham, Frances Jourd’heuil, James G. Traylor, A Wayne Orr, David Jourd’heuil

## Abstract

Rupture of the fibrous cap is the primary cause of clinical complications from atherosclerosis. Smooth muscle cells (SMCs) are a major contributor to fibrous cap development and stability through de-differentiation to extracellular matrix-producing cells. We previously showed that the antioxidant enzyme cytoglobin (CYGB) is expressed in vascular SMCs and regulates SMC dependent vascular remodeling and gene expression. In the present study, we investigated the function of SMC-CYGB in atherosclerosis. To this end, we generated a mouse line with SMC-specific deletion of *Cygb* and simultaneous SMC-lineage tracing. We found that SMC specific deletion of CYGB increased fibrous cap thickness in a 17-week PCSK9-AAV8 gain of function combined with Western diet mouse model of atherosclerosis. SMC specific deletion of CYGB increased collagen deposition and SMC cellularity of the fibrous cap in the absence of changes in total plaque and necrotic core sizes. CYGB expression in SMCs was associated with transdifferentiation towards a fibroblast-like, matrix remodeling phenotype. Finally, CYGB was expressed in the fibrous cap of human coronary atherosclerotic lesions and was associated with ACTA2 positive cells. These results provide first-time evidence that SMC-CYGB reduces plaque stability by decreasing cap thickness, collagen deposition, and SMC cellularity.

## Introduction

Despite advances in the diagnosis and treatment of atherosclerosis, acute coronary disease due to the rupture of unstable atherosclerotic plaques and expulsion of prothrombic contents remains a leading cause of comorbidity and mortality worldwide^1–6^. Studies over the past two decades have established the importance of the fibrous cap that overlays the necrotic core of atherosclerotic lesions as a key determinant of lesion stability. Lesions with thick fibrous caps that are rich in extracellular matrix (ECM) and ACTA2 positive cells are less likely to rupture^2,7–11^ , and strategies aimed at promoting a stable fibrous cap may hold significant therapeutic potential.

The origin of ECM producing cells in the fibrous cap is heterogenous with smooth muscle derived cells (dSMCs) contributing to the largest fraction of cells^2,7–14^. Overall, lineage tracing studies in mouse models have established that pro-atherogenic signals stimulate the oligoclonal expansion and de-differentiation of medial SMCs to give rise to a variety of cells with different functional attributes including protective ECM producing fibroblasts and myofibroblast-like dSMCs^15–19^. Smooth muscle specific knockout studies (e.g. TCF21, KLF4, OCT4, IL-1β, PDGF, and MCP1^15,17,20–25^ ) have provided mechanistic insights into the phenotypic transition of VSMCs and determined that some dSMC subtypes also play detrimental roles. Combined with genome wide associations studies indicating that some loci that confer cardiovascular associated risks are associated with smooth muscle cells^26–28^, this indicates that dSMCs play an important role in the development of atherosclerosis, including the establishment of a fibrous cap.

At the cellular level, atherosclerosis is associated with increased oxidative stress that drives lesion progression^29–31^. Work over the past two decades also indicate an important role for redox signals in regulating vascular cell function in atherosclerosis ^30,32–36^. Accordingly, studies aimed at manipulating cellular sources of ROS in vascular SMCs have shown that redox signaling contributes to the phenotypic modulation of these cells ^32,37–41^. However, the exact role of these redox signals in vascular SMC phenotypic transition within the atherosclerotic plaque and their contribution to the development of a protective fibrous cap are less clear. Information on the role of specific antioxidant systems and how they may impact SMC redox signaling is also limited. For example, peroxiredoxins and glutaredoxins – two essential reducing systems for the removal of intracellular hydrogen peroxide – have been shown to regulate vascular remodeling^42–47^. However, their specific role in dSMC function in the atherosclerotic lesion is unknown.

We and others have shown that the antioxidant enzyme cytoglobin (gene code: CYGB) scavenges hydrogen peroxide and is expressed in mouse and human vessels in medial vascular SMCs and adventitial fibroblasts^48–57^. *In vitro*, we found that cytoglobin deficiency is associated with complex changes in SMC gene expression^48^. We also established that cytoglobin deficiency in two rodent models of vascular remodeling inhibits neointima formation^49^. The present study investigated whether cytoglobin deficiency in SMCs exacerbates atherosclerosis. For this purpose, we generated a mouse line with inducible SMC specific deletion of CYGB combined with SMC lineage tracing. Hypercholesterolemia was induced with an AAV8-PCSK9 virus and Western diet to promote atherosclerotic lesion development. While we found no overall change in plaque size, our results indicate that SMC-cytoglobin decreases the contribution of SMCs to the fibrous cap and ultimately decreases cap thickness. These results are consistent with a significant role for cytoglobin in regulating vascular SMC fate and plaque stability.

## Methods

### Analysis of Single Cell RNA-Seq Data

All analyses and visualizations were performed using publicly available data. Single-cell RNA-sequencing data were obtained from the gene expression omnibus under accession number GSE155514 (mouse) and GSE159677 (human). Processed expression matrices were analyzed using Seurat version 5.4.0 in R version 4.4.2^58^. Cells and genes were filtered based on standard quality control metrics (e.g., gene count, mitochondrial content) as described in the original publications^18,59^. For the mouse dataset, SMC lineage was determined by Authors FACS sorting as previously described^18^. Dimension reduction was performed and plotted with Seurat using uniform manifold approximation and projection (UMAP)^58^. Potential of heat-diffusion for affinity-based trajectory embedding (PHATE) were generated using the packages PHATE, MAGIC, and plotted with ggplot2 ^60–62^. Differential gene expression analysis was conducted between cell populations using Seurat default parameters, and upregulated genes were defined based on threshold > 1.5, adjusted P value < 0.05. Gene ontology enrichment analysis was performed using the clusterProfiler R package^63^, with the genome annotation databases org.Mm.eg.db for mouse data, and org.Hs.eg.db for human data^64,65^.

### Mouse Line and Procedures

All animal protocols were approved by the Institutional Animal Care and Use Committee at Albany Medical College. Mice were housed under standard laboratory conditions with a light cycle of 12 hours, and had free access to water as well as standard chow diet (22% kCal fat, 23% kCal protein, 55% kCal Carbohydrates; PicoLab, Cat# 5058) or “Western” purified atherogenic diet (42% kCal Fat, 15.2% kCal Protein, 42.7% kCal Carbohydrates; Inotiv, Cat# 88137) for atherosclerosis experimentation. Mice were bred in-house in Albany Medical College Animal Resources Facility. Myh11-CreER^T2^ (CRE^+/0^), which were a kind gift from Dr. Singer (Albany Medical College), were crossed with B6.Cg-Gt(ROSA)26Sor^tm6(CAG-ZsGreen1)Hze^/J (Ai6(RCL-ZsGreen) (ZsGreen^+/+^) obtained from Jackson laboratories (JAX:007906) to generate mice that were ZsGreen^+/+^,Cre^+/0^. The resulting mice are crossed with B6NCrl-*Cygb*^tm1c(EUCOMM)Wtsi^ (TM1C^+/+^) obtained from the University of Toronto Canada to generate TM1C^+/-^, ZsGreen^+/+^, Cre^+/0^ mice. These TM1C heterozygous mouse pairs were bred to generate TM1C^+/+^, ZsGreen^+/+^, Cre^+/0^ (Cygb^SMC^(-/-)) CYGB floxed and TM1C^-/-^, ZsGreen^+/+^, Cre^+/0^ (Cygb^SMC^(+/+)) CYGB wild type littermate controls. Genomic DNA was extracted from ear clips, and genotyping was performed using PCR amplification with the following primers: Cygb wild type: sense, 5’-TTGGTCCACACCCGCTTCTTCATCTG-3’; Cygb wild type: antisense, 5’-TCGCTGAGACATTAGAGGAGCTCACAG-3’; Cygb flox (TM1C): sense, 5’-TCAGCATCCAAATGGGTCCCTGTCC-3’; Cygb flox (TM1C): antisense, 5’-TGCTATACGAAGTTATCTCGACGAAGTTCC-3’; ZsGreen1 wild type: sense, 5’-AAGGGAGCTGCAGTGGAGTA-3’; ZsGreen1 wild type: antisense, 5’-CGAAAATCTGTGGGAAGTC -3’; ZsGreen1 transgene: sense, 5’-AACCAGAAGTGGCACCTGAC-3’; ZsGreen1 transgene: antisense, 5’-GGCATTAAAGCAGCGTATCC-3’; Cre transgene: sense, 5’-TCCAACCTGCTGACTGTG-3’; Cre transgene: antisense, 5’-TCAGAGTTCTCCATCAGGG-3’. The Myh11-CreER^T2^ is located on the Y chromosome limiting experimentation to male individuals. For linage tracing and CYGB deletion studies, genetic recombination was initiated in 7 week old littermates by intraperitoneal injection of 2mg of tamoxifen for 5 days to activate Cre-recombinase and allowed 9 days to excrete excess tamoxifen. For the atherosclerosis model, 9 week old mice were subjected to a single retro-orbital injection with 100µl of 1.75 x 10^9^ pc/µl AAV8-D377Y-mPCSK9 gain of function virus(Vector Biolabs, SKU VB-377Y), and diet provided is switched to “Western” diet. Three weeks post injection, plasma cholesterol levels were assessed using a cholesterol E kit (Fujifilm Healthcare Solutions, Cat# 999-02601 ). Mice with plasma cholesterol levels > 1000 mg/dL were selected for inclusion in subsequent atherosclerosis experiments. 16 to17 weeks post injection, mice were fasted for 6-8 hours, and blood glucose was measured using an Accu-Chek Aviva Plus glucometer. At 17 weeks post injection, mice were euthanized by CO_2_ asphyxiation and then perfused with saline into the left ventricle.

### Murine Tissue Preparation for Lesion Morphology and Staining

The hearts of the mice were fixed in 10% neutral buffered formalin (NBF) (Sigma, HT501320) and cryopreserved in 30% sucrose solution for 24-48hrs, and frozen in Optimal Cutting Temperature Medium (OCT) (Sakura, REF: 4583) buffered by 2-methyl butane (Sigma, Cat# M32631). Sections were cut 10µm thick using a Leica CM1850 cryostat and transferred onto charged microscope slides (Electron Microscopy Sciences, Cat# 63700W1). Slides were then dried to room temperature and stored at -80°C until use.

### Murine Immunofluorescence Staining

Thawed, dried mouse heart sections were subjected to antigen retrieval in 100°C 10 mM citrate buffer (Sigma, Cat# C9999) for 30 minutes, after cooling to room temperature the slides were washed in Phosphate Buffered Saline with 0.1% Triton X 100 (PBST). Sections were then blocked with 5% serum representing the secondary antibody species, 1% bovine serum albumin, dissolved in PBST for one hour at room temperature, then incubated with the following primary antibodies or controls overnight at 4°C: Cytoglobin Rabbit pAB: Sigma, Cat# HPA017757, Alpha Smooth Muscle Actin mAB (Acta2): Invitrogen, Cat# MA5-11547, Mouse IgG isotype control: Vector labs, Cat# I-2000, Rabbit IgG isotype Control: Vector labs, Cat# I-1000. After three rinses in PBST each section was incubated with the following fluorescent conjugated secondary antibody for one hour at room temperature: Goat anti-Rabbit IgG: Invitrogen, Cat# A32740, Goat anti-Mouse IgG: Invitrogen, Cat# A21236. Following three washes in PBST the nuclei were counterstained with 1 µM 4′,6-diamidino-2-phenylindole (DAPI) (Sigma, Cat# D9542) for 15 minutes at room temperature. After a final rinse coverslips were mounted on each slide with VectaShield Vibrance Antifade Mounting Medium (Vector labs, Cat# H-1700), sealing with clear nail polish and the slides were stored protected from light at 4°C until imaging. Images for mouse sections were acquired using a Leica Thunder microscope using the manufacturer’s software under identical acquisition settings. Cell counting using Cytation 5 Gen 5 software.

### Hematoxylin and Eosin (H&E) Staining

Slides with 10 µm OCT embedded tissue were removed from the -80°C freezer and immediately immersed in a fixative solution including 70% ethanol, 4% formaldehyde, 5% glacial acidic acid for 3 minutes. The sections were dehydrated with 70% and 90% ethanol for one minute each, washed with tap water then rinsed in deionized water for 30 seconds. Nuclear staining was achieved with a three-minute incubation in Hematoxylin (Fisher Scientific, Cat # 22-050-111). Excess dye was removed with flowing tap water before clarification and bluing. After a quick rinse in 70% ethanol the cytoplasmic and extracellular components were counterstained with EosinY (Fisher Scientific, Cat# 22-050-110) for 30 seconds. Tissues were rinsed and dehydrated with increasing concentrations of ethanol to 100%, cleared with xylenes (Sigma, Cat# 534056), and coverslipped with acrylic resin mounting media. Tissue sections are visualized on an Olympus BX51 microscope using cellSens standard software (version 4.2.1).

### Masson Trichrome Staining

OCT embedded sections were fixed again in 10% NBF for 1 hour. Sections were incubated in Boin’s solution (Polysciences, Cat# 25088A) overnight at room temperature and rinsed in tap water to remove picric acid. Sections were incubated in Weigert’s Iron Hematoxylin (Polysciences, Cat# 25088B1 & 25088B2) working solution for 10 minutes followed by a distilled water rinse to remove unbound dye. Sections were then incubated in Biebrich Scarlet with Acid Fuchsin solution (Polysciences, Cat# 25088C) for 5 minutes to visualize the muscle and cytoplasm, rinsed in distilled water, then differentiated in Phosphomolybdic/Phosphotungstic acid solution (Polysciences, Cat# 25088D) for 10 minutes. Sections were then incubated in Aniline Blue solution (Polysciences, Cat# 25088E) for 5 minutes to mark the collagen fibers, followed by a one-minute incubation in 1% Acetic Acid solution (Polysciences, Cat# 25088F). Followed by a final rinse sections were dehydrated in increasing concentrations of ethanol to 100% for 2 minutes each and cleared in xylene. Slides were coverslipped with acrylic resin mounting medium prior to imaging on a NanoZoomer 2.0-RS slide scanner (Hamamatsu).

### Human Immunofluorescent Staining

Human coronary artery tissue was utilized from the Louisiana Coronary Artery (LOCATE) biobank through a collaboration with Dr. Wayne Orr (LSU Health Shreveport). Deemed non-human research by the local IRB due to the exclusive use of post-mortem tissue (STUDY00001712), the LOCATE Biobank includes over 1000 human coronary arteries (right coronary artery, left anterior descending artery, circumflex artery) excised post-mortem during routine autopsy from over 500 patients. Plaques were preserved in 4% formaldehyde, embedded in paraffin, and cut into 5 µm sections prior to staining.

Sections were deparaffinized and unmasked by heating in Trilogy buffer (Sigma-Aldrich; cat 920P) using the high cycle on a pressure cooker. After a wash in PBS the sections were permeabilized/quenched in PBS buffer containing 0.1% Triton-X 100 and 0.1M Glycine for 5 minutes at room temperature. Slides were then blocked with 5% serum representing the secondary antibody species, 1% bovine serum albumin, dissolved in PBST for one hour at room temperature. This was followed with incubation with the following primary antibodies or controls overnight at 4°C: Cytoglobin Rabbit pAB: Protein Tech, Cat# 13317-1-AP, Alpha Smooth Muscle Actin mAB (Acta2): Invitrogen, Cat# MA5-11547, Mouse IgG isotype control: Vector labs, Cat# I-2000, Rabbit IgG isotype Control: Vector labs, Cat# I-1000. Unbound primary antibodies were washed away in PBST prior to a one-hour incubation with secondary antibody incubation for one hour at room temperature: Goat anti-Rabbit IgG: Invitrogen, Cat# A32740, Goat anti-Mouse IgG: Invitrogen, Cat# A21236. Nuclei were counterstained with 1 µM DAPI for 15 minutes at room temperature, following a brief wash each slide was coverslipped with VectaShield Vibrance Antifade Mounting Medium (Vector labs, Cat# H-1700). Images were acquired using a BioTek Cytation 5 multimodal imaging system (Agilent 169 Technologies, Inc.). Images were processed in Image J.

### Statistical Analysis

All statistical analysis were performed using GraphPad Prism version 9.0.0. Data is represented as mean ± SEM. Unpaired Student’s t-tests were used to compare lesion area, percent necrosis, percent collagen, fibrous cap thickness, cell counts. Statistical Significance indicated by *p < 0.05, **p< 0.01. Sample sizes are indicated in the figure legends. Lesion area and necrotic area analyses were performed on H&E-stained lesion cross-sections using ImageJ software. Fibrous cap thickness and collagen positive area analyses were performed on ACTA2 immunofluorescent-stained, or Masson Trichrome-stained lesion cross-sections. Average fibrous cap thickness was calculated as ∑area (um^2^) divided by ∑shortest longest distance (µm), using the skeletonize function in ImageJ.

### Reagents List

Additional reagents list found in **Supplemental Table 2**.

## Results

### Cytoglobin-Expressing Smooth Muscle Derived Cells Acquire Fibroblast-Like Features in the Atherosclerotic Plaque

We first determined the cellular distribution of CYGB in atherosclerosis. To this end, we reanalyzed a publicly available single cell dataset (GSE155514) obtained from mouse aortas, in which SMCs were lineage-traced for ZsGreen1 expression, and samples obtained at increasing duration of atherosclerosis induction (0, 8, 16 and 26 weeks)(**Fig. 1A, Supplemental Fig. 1A**)^18^. Following hierarchical clustering, we identified 17 unique cellular clusters, with 9 cell types (**Fig. 1A, Supplemental Fig. 1B, C**). We found that CYGB mRNA transcripts were enriched in fibroblast clusters originating from dSMCs (ZsGreen1 positive; **Fig. 1B**). In contrast, dSMC-CYGB transcripts were only minimally observed in other related dSMC clusters (**Fig.1B, Supplemental Fig. 1F, G**).

**Figure 1:**
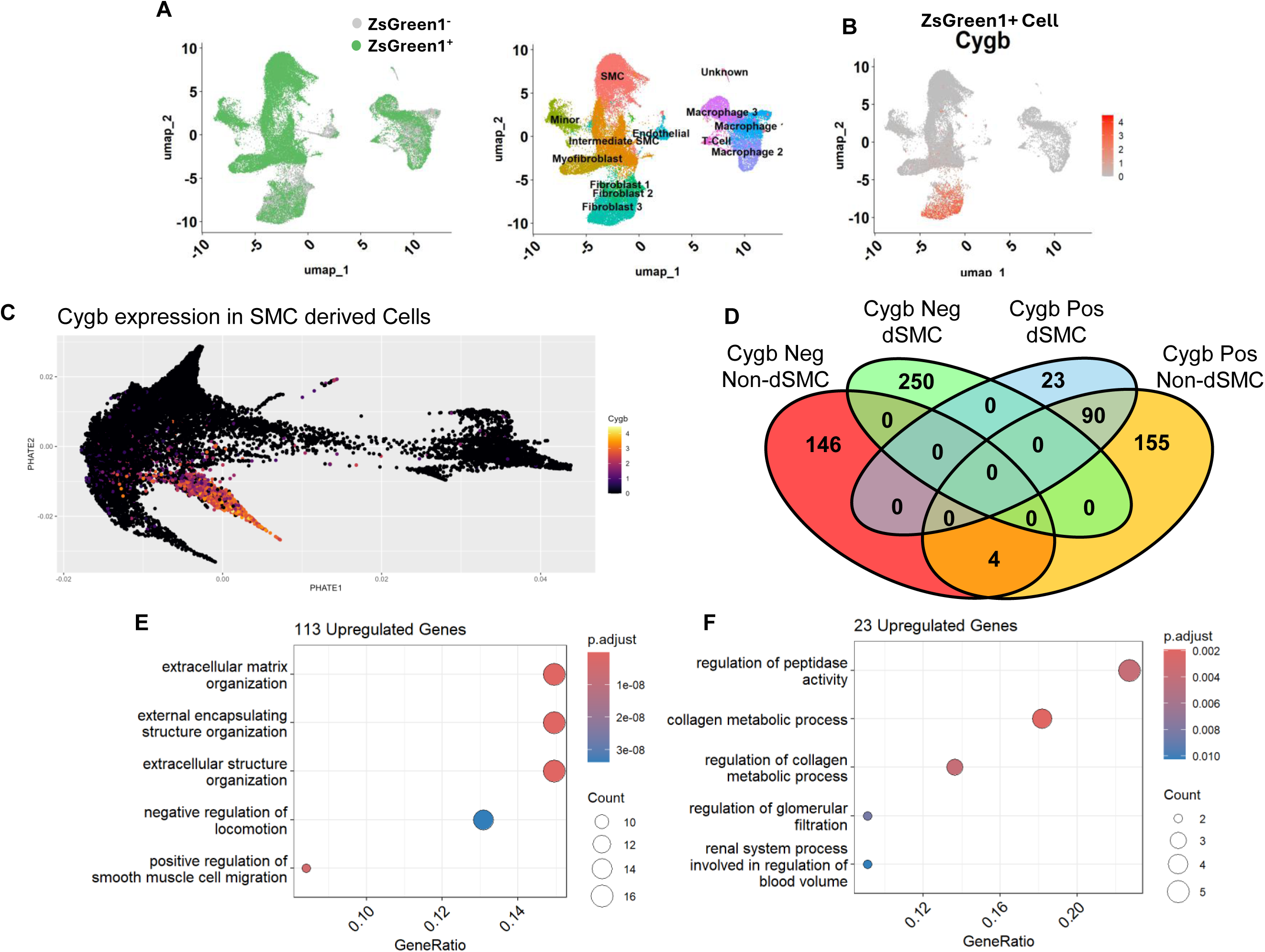
Single Cell RNA Seq dataset from Publicly Available Dataset Identifies Cygb-Expressing SMCs Have a Distinct Transcription Profile (GSE155514). **A)** Integrated UMAP projections identifying ZsGreen1^+^ and ZsGreen1^-^ cells, and cluster identity from a publicly available scRNA-seq dataset. **B)** Feature plot of ZsGreen1^+^ cells for the corresponding gene transcript expression level indicated by color scale. **C)** Phate projection of ZsGreen1^+^ cells for the mRNA expression of Cygb. **D)** Venn diagram illustrates the overlap of differentially expressed upregulated genes between CYGB and ZsGreen1 conditions. **E)** Top five gene ontology terms assosiated exclusively with CYGB^+^ ZsGreen1^+^ cells. **F)** Top five gene ontology terms of all genes assosiated with CYGB^+^ ZsGreen1^+^ cells. Adjusted p values < .05, average log_2_ expression > 1.5.

In agreement with CYGB expressed in fibroblast clusters of dSMCs, PHATE projection of dSMCs indicated that CYGB expression was associated with a specific differentiation trajectory of dSMCs in atherosclerosis (**Fig. 1C**). To further understand the functions of cells expressing CYGB, we performed differential gene expression analysis comparing dSMCs and non-SMCs with and without CYGB expression. A total of 113 genes were upregulated in the CYGB-positive dSMCs (**Fig. 1D**). Gene ontology analysis for these 113 genes included “extracellular matrix organization”, “collagen formation and metabolic processes”, and “negative regulation of locomotion” pathways (**Fig. 1E**). Of the 113, only 23 of these genes were specific to the CYGB-positive dSMCs. These 23 genes were associated with gene ontology terms including “collagen metabolic process”, “regulation of collagen metabolic process”, and “regulation of peptidase activity” pathways (**Fig. 1F**). Overall, these results suggest that CYGB expression in dSMCs is associated with fibroblast transitioned cells with enriched matrix remodeling activity.

### SMC Specific Deletion of Cytoglobin Increases Fibrous Cap Thickness and Increases SMC Derived Cell Content of the Fibrous Cap

Next, we determined the effect of deleting CYGB specifically in SMCs on the development of atherosclerotic plaques. During transdifferentiation, SMCs lose many of their canonical markers making it difficult to properly identify these cells in complex disease environments such as atherosclerosis . To avoid this issue, we generated a new mouse line allowing for the lineage tracing of SMCs, while permitting the specific deletion of CYGB. We first crossed transgenic mice, which express tamoxifen-inducible Cre-recombinase driven by the SMC-specific Myh11 promoter with a Cre-responsive ZsGreen1 inserted at the ROSA26 locus (**Fig. 2A**). The resulting mice were then crossed with a novel floxed CYGB mouse line, allowing for the same Myh11 promoted Cre system to specifically genetically delete CYGB in SMCs upon tamoxifen activation (**Fig. 2A**). To confirm SMC-specific loss of CYGB, we immuno-stained for CYGB in the descending thoracic aorta of lineage traced CYGB wild type (Cygb^SMC^(+/+)) and lineage traced CYGB smooth muscle specific knockout (Cygb^SMC^(-/-)) mice (**Fig. 2B**). We confirmed the specific loss of CYGB protein expression in the ZsGreen1 positive cells of Cygb^SMC^(-/-) mice, while the expression of CYGB in non-SMCs persisted in the adventitia (**Fig. 2B**).

**Figure 2:**
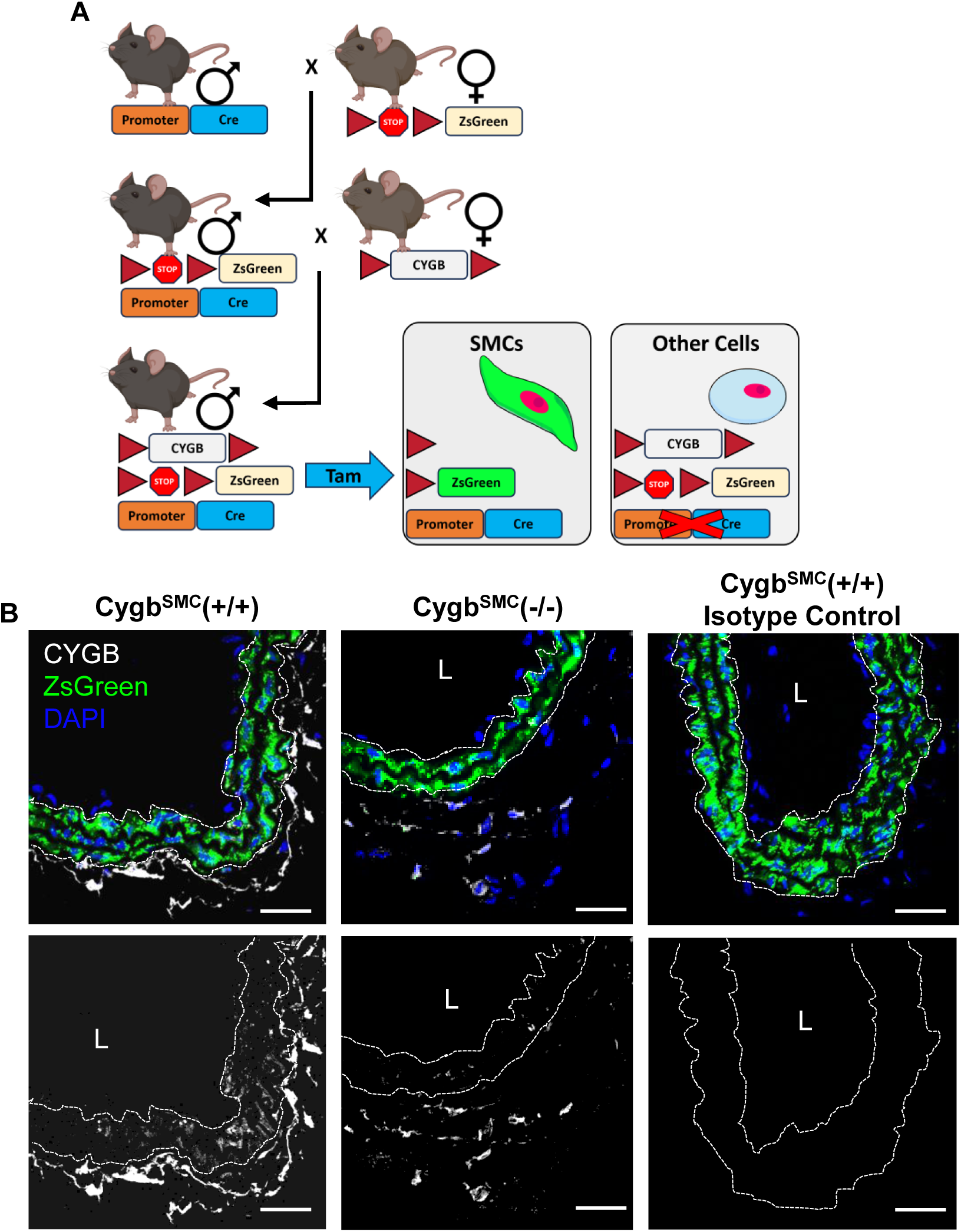
Establishment of SMC Specific Cygb Knockout with Lineage Tracing. **A**) Illustration of breeding scheme used to generate conditional lineage tracing and conditional genetic knockout mice. **B)** Representative left common carotid artery sections of CYGB^SMC^(+/+) and CYGB^SMC^(-/-) mice, immunofluorescent stained for Cytoglobin (white) with endogenous ZsGreen1(green) flourecence and nuclei counterstained with DAPI (blue). L = Lumen, dashed lines show medial layer delineated by ZsGreen1 expression, scale bar represents 50μm.

To induce plaque development, we injected Cygb^SMC^(+/+) and Cygb^SMC^(-/-) mice with an AAV8 gain of function PCSK9 (AAV8-GOF-PCSK9) virus and provided a high fat western diet for 17 weeks (**Fig. 3A**). Importantly, there was no statistical difference in plasma cholesterol, or blood glucose between groups (**Supplemental Fig. 2A, B**). We confirmed that CYGB was expressed within lesion dSMCs of the of Cygb^SMC^(+/+) mice and lost in dSMCs of Cygb^SMC^(-/-) mice (**Fig. 3B**). Interestingly, we found positive staining for CYGB in cells that lacked the expression of ZsGreen1 in both the Cygb^SMC^(+/+) and Cygb^SMC^(-/-) mouse groups (**Fig. 3B**), suggesting expression of CYGB in non-dSMCs. This was surprising, as our analysis of the non-dSMC populations indicate the expression of CYGB is primarily localized to the fibroblast clusters, which are not typically associated with atherosclerotic lesion formation (**Supplemental Fig. 1E,F**).

**Figure 3:**
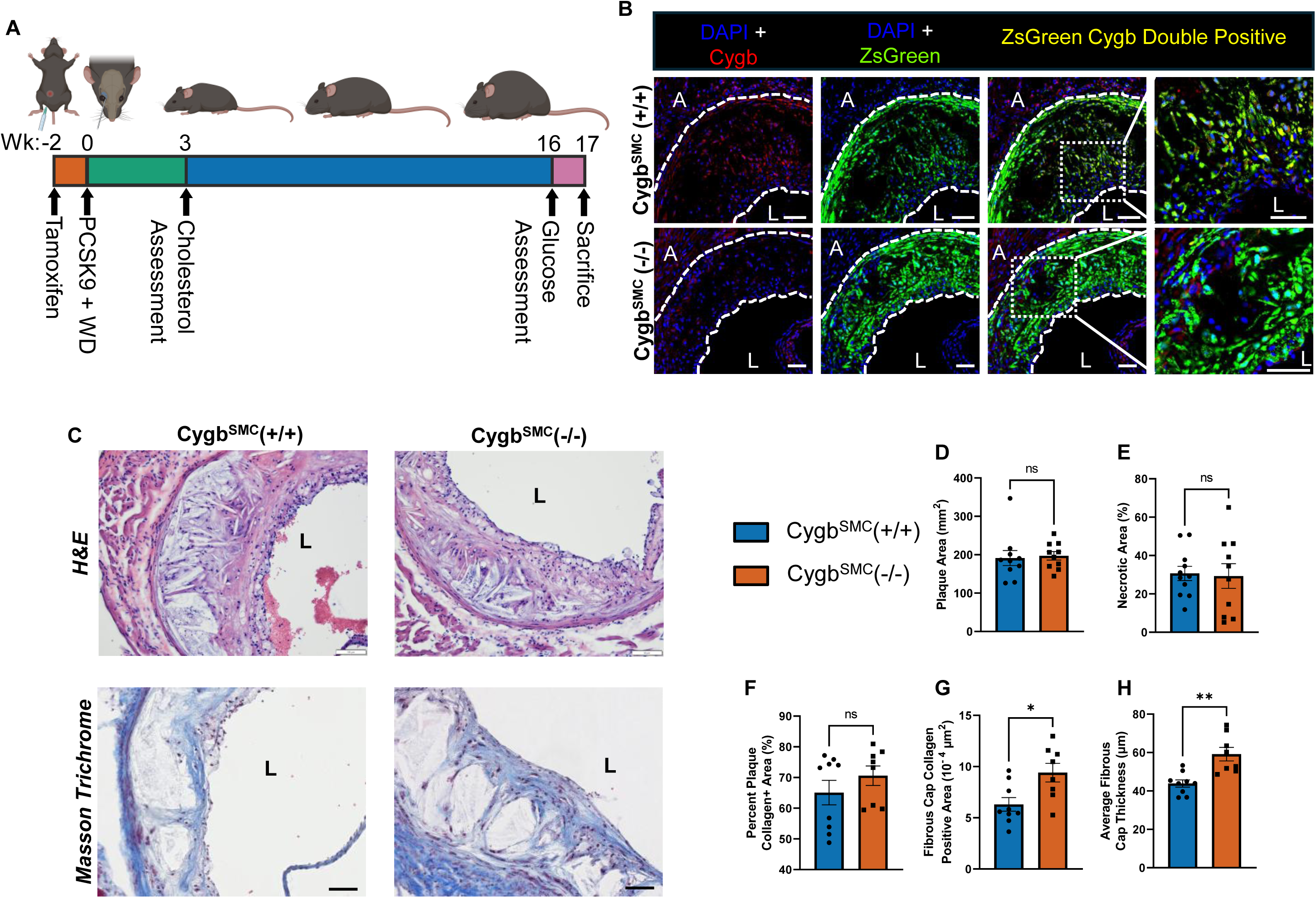
SMC Specific Deletion of Cygb Increases Fibrous Cap Thickness Without Altering Lesion Size. **A)** Schematic overview of the treatment regimen of mice. **B)**Immunofluorescence staining for DAPI (blue) and Cygb (red) with endogenous ZsGreen1 (green) of plaques from the aortic sinus of Cygb^SMC^(+/+) and Cygb^SMC^(-/-). L = Lumen, scale bar represents 50μm, dashed lines indicate medial layer and plaque. **C)** Plaque aortic sinus sections of mouse groups, stained for Hematoxylin and Eosin (H&E) (top) and Masson Trichrome (bottom). **D)** Quantification of plaque area within aortic sinus (mm^2^). **E)** Percentage of plaque area classified necrotic (%). **F)** Percent of plaque area which is collagen positive (%). **G)** Collagen positive area within the fibrous cap (10^-4^ μm^2^). **H)** Average Fibrous cap thickness (µm). All results are represented as mean ± SEM and each dot represent an individual mouse. Student’s t-test was used. ns p> .05, * p< 0.05, ** p< 0.01, *** p< 0.001

Next, we performed H&E and Mason’s trichrome staining to determine the effect of SMC-specific deletion of CYGB on plaque morphology (**Fig. 3C**). Quantification of these stains revealed no difference in total plaque area, necrotic core area, or total collagen content between Cygb^SMC^(+/+) and Cygb^SMC^(-/-) mice (**Fig. 3D-F**). However, there was a significant increase in the collagen positive area of the fibrous cap and an increase of the fibrous cap thickness in Cygb^SMC^(-/-) mice compared to the control mice (**Fig. 3G,H**). Overall, these results suggest that SMC-CYGB may influence plaque stability by regulating fibrous cap thickness and collagen content.

### Smooth Muscle Specific Deletion of Cytoglobin Increases SMC Derived Cell Content of the Fibrous Cap

For the immunofluorescence staining studies, we used ACTA2 staining to identify the cap region of the lesions. First, we showed strong expression of CYGB within the fibrous cap of lesions in Cygb^SMC^(+/+) mice (**Fig. 4A**), and co-association of CYGB with ACTA2 was evident in both ZsGreen1^+^ dSMCs and ZsGreen1^–^ non dSMCs (**Fig. 4A**). Consistent with **Fig 3E**, fibrous cap thickness based on ACTA2 coverage was significantly increased in Cygb^SMC^(-/-) mice (**Fig. 4B**). We also found that the number of ZsGreen1^+^ cells increased within the fibrous caps of Cygb^SMC^(-/-) mice (**Fig. 4C**), even when expressed relative to cap area to account for differences in fibrous cap size (**Fig. 4D**). Overall, these results indicate that loss of SMC-CYGB increases the fraction of dSMCs occupying the cap, suggesting a critical role for SMC-CYGB in regulating fibrous cap cellular composition and cap thickness.

**Figure 4:**
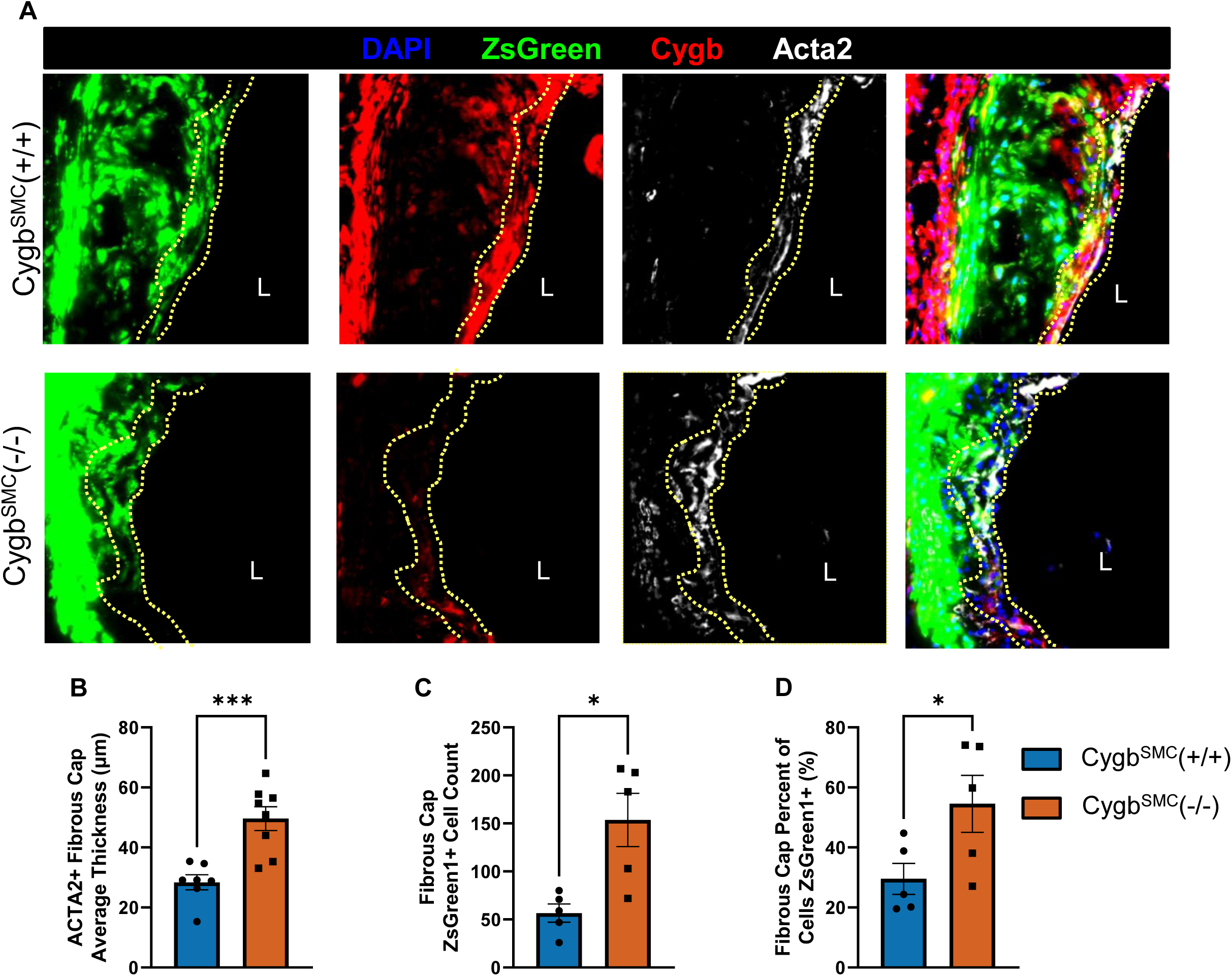
Smooth muscle specific genetic deletion of cytoglobin increases SMC derived cellularity of the fibrous cap. **A)** Immunofluorescence staining for CYGB (red), ACTA2 (white), counterstained with DAPI (blue), and endogenous ZsGreen1 (green) of plaques from the aortic sinus of Cygb^SMC^(+/+) and Cygb^SMC^(-/-) mice. L = Lumen. Dashed lines show fibrous cap delineated by Acta2 expression. **B)** Average fibrous cap thickness determined by Acta2 area (μm). **C**) Total number of ZsGreen1 positive cells within the Acta2 positive fibrous cap area. **D)** Percent of cells positive for ZsGreen1 within the Acta2 positive region of the fibrous cap (%). All results are represented as mean ± SEM and each dot represent an individual mouse. Student’s t-test was used. ns p> .05, * p< 0.05, ** p< 0.01, *** p< 0.001

### Cytoglobin is Expressed in ACTA2 Positive Cells in the Fibrous Cap of Human Coronary Atherosclerotic Lesions

To assess CYGB expression in human atherosclerotic plaques, we reanalyzed a publicly available scRNA-seq dataset (GSE159677), comprising cells isolated from the atherosclerotic core and proximal adjacent regions of human atherosclerotic plaques (**Fig. 5A**). We identified 20 subclusters within this dataset, with 7 distinct cell types (**Fig. 5B, 5C, Supplemental Fig. 3A**). CYGB expression was enriched within the SMC and endothelial cell populations of the atherosclerotic core and proximal adjacent region (**Fig. 5D**). Notably, significant enrichment of CYGB transcripts was observed within cluster 19 SMCs of the proximal adjacent region (**Fig. 5E**). Gene ontology analysis of the dSMC clusters comparing CYGB^+^ dSMCs and CYGB^-^ dSMCs indicated enrichment of pathways related to negative regulation of cell migration and cell adhesion (**Fig. 5F**), while pathways associated with muscle development and contraction were downregulated (**Fig. 5G**).

**Figure 5:**
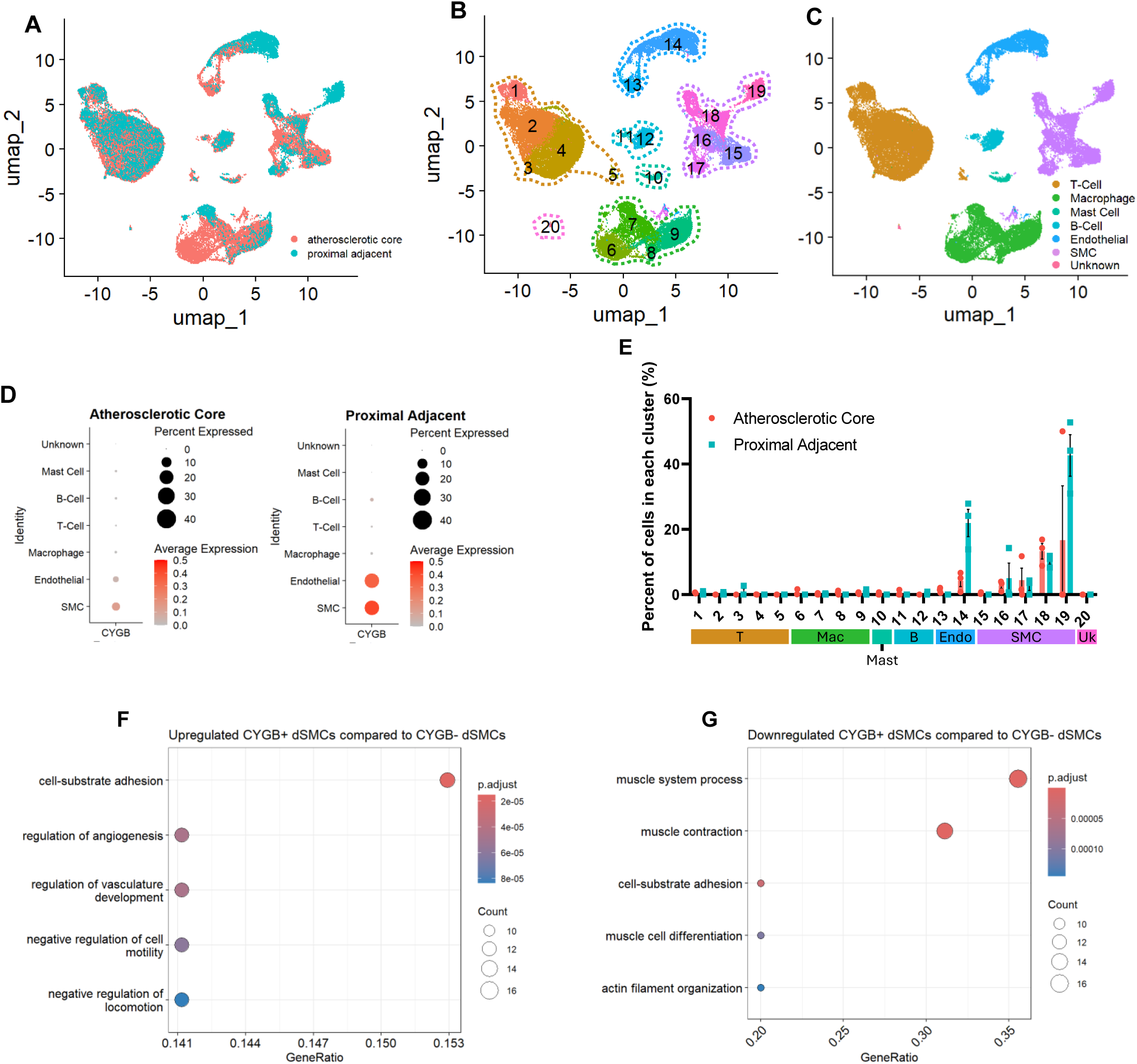
Validation in human atherosclerosis. A,B,C) UMAP projection of human atherosclerosis explants from publicly available scRNA-seq data set (GSE159677) identifying anatomical location (A), cell cluster (B), and cell type (C). **D)** Feature plot identifying cytoglobin mRNA expression. **E)** Quantification of the frequency of cytoglobin positive cells in each cluster as a percentage of total cells in each cluster. **F)** Differential gene expression between up- and down-regulated genes between cytoglobin positive and negative smooth muscle derived cells. Average log2 threshold ± 1.5, adjusted p value < .05.

Finally, we immuno-stained tissue sections obtained from atherosclerotic human coronary arteries (**Supplementary Table 1**). We found CYGB staining within the fibrous caps of the lesions in addition to the adventitia, media, and intimal layers (**Fig. 6A**). Importantly, there was strong co-association of CYGB with ACTA2 positive cells in the caps of these samples (**Fig. 6A**).

**Figure 6:**
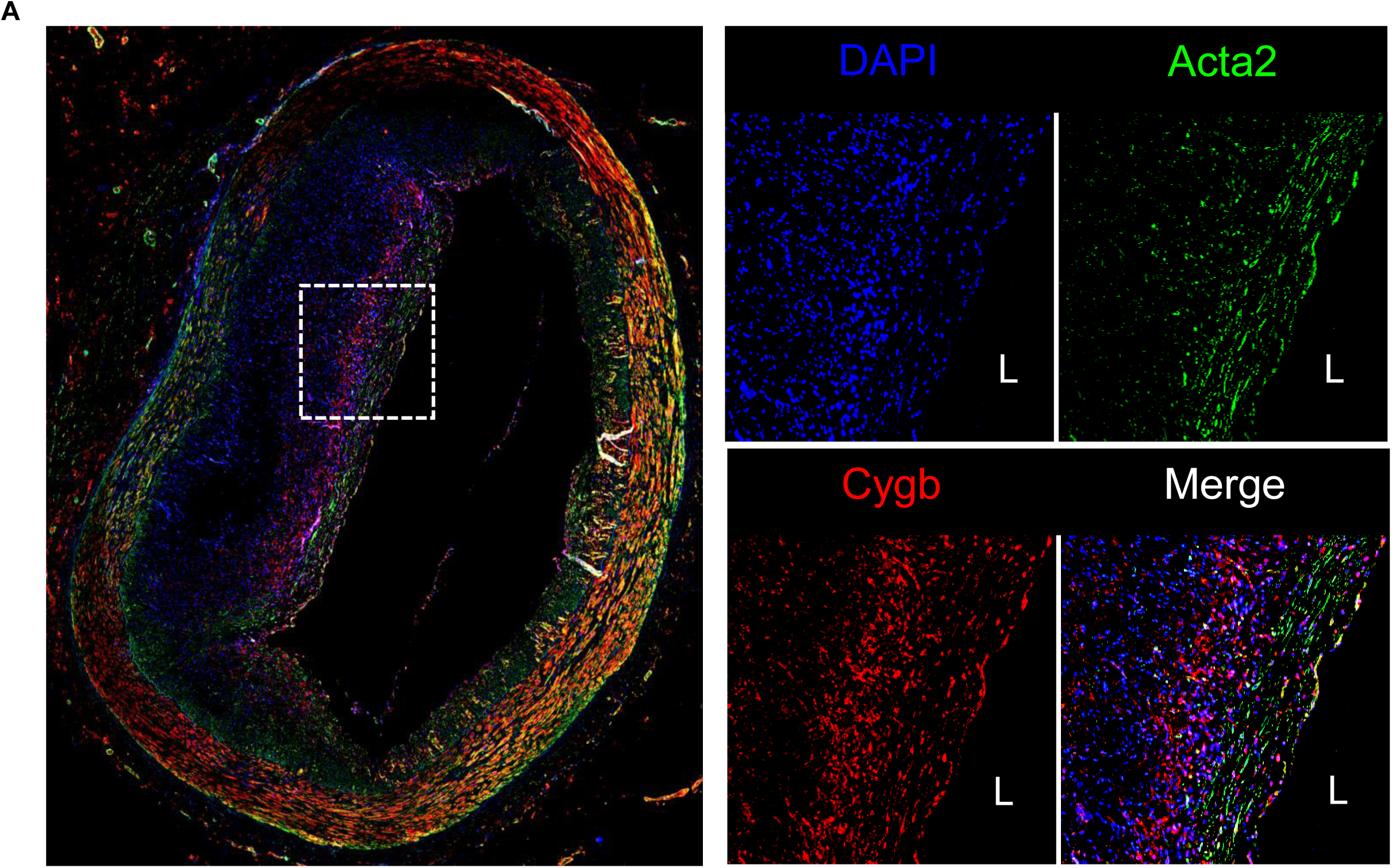
Human Atherosclerotic plaques contain Cytoglobin. **A)** Representative images of human atherosclerotic plaque stained for Acta2 (green), Cytoglobin (red), and counterstained with DAPI (blue).

## Discussion

In the present work, we established the role of cytoglobin (CYGB) in regulating smooth muscle cell (SMC) function in atherosclerosis. We chose to study the role of CYGB because we previously showed that this antioxidant enzyme plays a key role in regulating smooth muscle-mediated hyperplasia *in vivo* after balloon angioplasty and carotid ligation injury^49^. In the present study, we found that SMC-specific deletion of CYGB increased the thickness and collagen content of the fibrous cap within atherosclerotic lesions using a 17-week PCSK9-GOF mouse model of atherosclerosis. This was accompanied by an increase in the number of smooth muscle derived cells (dSMCs) within the fibrous cap. We also established that CYGB is primarily expressed in atherosclerotic lesions within dSMCs that have acquired fibroblast-like phenotypes. Finally, we show that CYGB co-associates with the fibrous cap of human atherosclerotic lesions. Overall, our results indicate for the first time that SMC CYGB is a negative regulator of plaque stability.

We had previously established that loss of CYGB inhibits neointima formation in the rat carotid angioplasty and mouse carotid ligation models^49^. Neointimal hyperplasia in these settings is the result of the de-differentiation, migration, and clonal expansion of medial SMCs to form these lesions ^67–74^. Thus, we posited earlier that CYGB deletion limits the capacity of SMCs to survive or undergo the phenotypic reprograming required for neointimal expansion^48,49^. Yet, the results in the current study indicate an increase in the number of dSMCs in the cap lesion and an increase in cap collagen content and cap thickness following SMC-specific deletion of CYGB. There might be several explanations for these discrepancies. In the first two studies, we performed targeted viral-mediated deletion of CYGB in the rat model of carotid angioplasty^49^ and global genetic deletion in the carotid ligation mouse model^48,49^. In addition to vascular SMCs, CYGB is expressed in adventitial fibroblasts in vessels. Although there is no evidence of CYGB expression in other cell types that might contribute to vascular remodeling in these two models, it is possible that the inhibition of neointima formation observed previously is a consequence of decreased CYGB in adventitial fibroblasts, in addition to SMCs or to more global effects associated with the global deletion of CYGB.

Another possibility is related to our current understanding of the contribution of SMCs to plaque formation and differences in SMC reprogramming between the different models. Indeed, past studies have established that de-differentiated vascular SMCs acquire distinct pathogenic states, which drive different aspects of plaque development. For example, ACTA2^+^ dSMCs contribute to the formation of a protective, collagen rich-fibrous cap that covers plaque lesions. Vascular SMCs may also transdifferentiate into myofibroblasts (FMCs) and chondromyocytes (CMCs) that promote plaque progressions. Notably, Amrute et al. recently identified a dSMC niche positive for fibroblast activation protein (FAP), which was comprised of FMCs and CMCs that could be targeted through immunotherapy to reduce plaque burden^75^. Importantly, our analysis indicates that CYGB is not expressed across all dSMCs but identifies fibroblast-like dSMCs in lesions. Overall, this suggests that SMC-specific deletion of CYGB alters the function of these fibroblast-like dSMCs and favors a fibrous cap with a greater content in ACTA2^+^ dSMCs. Although abundant within the atherosclerotic lesion, the role of fibroblast-like dSMCs is not yet fully understood^9,17,18,76^. Deciphering the function of these SMC subsets and specific mechanistic attributes such as CYGB-dependent pathways will require additional studies including single cell resolution approaches and possibly dual recombinase systems.

Studies from this laboratory and others have established that CYGB exerts strong antioxidant functions including hydrogen peroxide scavenging^48,50–52^. In subcultured human primary SMCs, CYGB regulate the cellular response to oxidative stress through nuclear interaction with the chromatin remodeler HMGB2 and alteration in DNA damage response and gene expression^48^. A study by Ou et al. indicates that treatment with recombinant CYGB is atheroprotective in a hyperlipidemic rat model^66^. These results most likely reflect the strong antioxidant activity of CYGB, in line with the well-characterized role of oxidative stress in atherosclerosis. In combination with our results showing strong expression of CYGB in ACTA2^+^ cells in human plaques, they highlight the therapeutic potential of CYGB in cardiovascular diseases. Determining the specific mechanisms by which CYGB may alter the function of vascular cells including SMCs and fibroblasts would therefore be essential for therapeutic applications.

## Acknowledgements

We would like to thank the laboratories of Dr. Gabrielle Fredman and Dr. Harold Singer for their assistance with retro-orbital delivery procedures.

## Supplemental Figure Legends

Supplemental Figure 1: **A)** Integrated UMAP projections identifying timepoints of western diet feeding from publicly available single cell dataset (GSE155514). **B)** UMAP visualizing 17 transcriptionally distinct cell clusters. **C)** Dot plot of traditional markers for cell identity. **D-E)** Dot plot showing CYGB expression by Western diet timeline in lineage traced ZsGreen1+ cells **(D)** and ZsGreen1-cells **(E)**. Dot size represents cells count, color represents percentage of cells expressing Cygb. **F)** Violin plots representing expression of Cygb between cell types in ZsGreen1+ and ZsGreen1- cells.

Supplemental Figure 2: **A)** Bar graphs showing mouse plasma cholesterol levels measured six hours after fasting and three weeks after PCSK9-AAV8-GOF virus and western diet (mg/dl). **B)** Bar graph showing plasma cholesterol 16 weeks after PCSK9-AAV8-GOF virus and western diet (mg/dL). Results are represented as mean ± SEM and each dot represent an individual mouse. Student’s t-test was used. ns p> .05, * p< 0.05, ** p< 0.01, *** p< 0.001

Supplemental Figure 3: **A)** Dot plot showing markers for identify cell identity.

## Abbreviations

CVD: Cardiovascular disease
CYGB: Cytoglobin
SMCs: Smooth Muscle Cells
dSMCs: Smooth Muscle Derived Cells
ECM: Extracellular Matrix
ROS: Reactive Oxygen Species
UMAP: uniform manifold approximation and projection.
PHATE: Potential of heat-diffusion for affinity-based trajectory embedding.

## References

1. Falk E, Shah PK, Fuster V: Coronary Plaque Disruption. Circulation, American Heart Association, 1995, 92:657–671.

2. Virmani R, Kolodgie FD, Burke AP, Farb A, Schwartz SM: Lessons From Sudden Coronary Death. Arteriosclerosis, Thrombosis, and Vascular Biology, American Heart Association, 2000, 20:1262–1275.

3. Naghavi M, Libby P, Falk E, Casscells SW, Litovsky S, Rumberger J, Badimon JJ, Stefanadis C, Moreno P, Pasterkamp G, Fayad Z, Stone PH, Waxman S, Raggi P, Madjid M, Zarrabi A, Burke A, Yuan C, Fitzgerald PJ, Siscovick DS, de Korte CL, Aikawa M, Juhani Airaksinen KE, Assmann G, Becker CR, Chesebro JH, Farb A, Galis ZS, Jackson C, Jang I-K, Koenig W, Lodder RA, March K, Demirovic J, Navab M, Priori SG, Rekhter MD, Bahr R, Grundy SM, Mehran R, Colombo A, Boerwinkle E, Ballantyne C, Insull W, Schwartz RS, Vogel R, Serruys PW, Hansson GK, Faxon DP, Kaul S, Drexler H, Greenland P, Muller JE, Virmani R, Ridker PM, Zipes DP, Shah PK, Willerson JT: From Vulnerable Plaque to Vulnerable Patient. Circulation, American Heart Association, 2003, 108:1664–1672.

4. Cardiovascular diseases (CVDs) [Internet]. [cited 2026 Mar 23], . Available from: https://www.who.int/news-room/fact-sheets/detail/cardiovascular-diseases-(cvds)

5. The top 10 causes of death [Internet]. [cited 2026 Mar 23], . Available from: https://www.who.int/news-room/fact-sheets/detail/the-top-10-causes-of-death

6. Ala-Korpela M: The culprit is the carrier, not the loads: cholesterol, triglycerides and apolipoprotein B in atherosclerosis and coronary heart disease. Int J Epidemiol 2019, 48:1389–1392.

7. Miano JM, Fisher EA, Majesky MW: Fate and State of Vascular Smooth Muscle Cells in Atherosclerosis. Circulation, American Heart Association, 2021, 143:2110–2116.

8. Allahverdianx S, Ortega C, Francis GA: Smooth Muscle Cell-Proteoglycan-Lipoprotein Interactions as Drivers of Atherosclerosis. In: von Eckardstein A, Binder CJ, editors. Prevention and Treatment of Atherosclerosis: Improving State-of-the-Art Management and Search for Novel Targets [Internet], Cham (CH), Springer, 2022 [cited 2026 Mar 23],. Available from: http://www.ncbi.nlm.nih.gov/books/NBK584292/

9. Bennett MR, Sinha S, Owens GK: Vascular smooth muscle cells in atherosclerosis. Circ Res 2016, 118:692–702.

10. Finn AV, Nakano M, Narula J, Kolodgie FD, Virmani R: Concept of Vulnerable/Unstable Plaque. Arteriosclerosis, Thrombosis, and Vascular Biology, American Heart Association, 2010, 30:1282–1292.

11. Bentzon JF, Otsuka F, Virmani R, Falk E: Mechanisms of Plaque Formation and Rupture. Circulation Research, American Heart Association, 2014, 114:1852–1866.

12. Shankman LS, Gomez D, Cherepanova OA, Salmon M, Alencar GF, Haskins RM, Swiatlowska P, Newman AAC, Greene ES, Straub AC, Isakson B, Randolph GJ, Owens GK: KLF4-dependent phenotypic modulation of smooth muscle cells has a key role in atherosclerotic plaque pathogenesis. Nat Med, Nature Publishing Group, 2015, 21:628–637.

13. Wang Y, Dubland JA, Allahverdian S, Asonye E, Sahin B, Jaw JE, Sin DD, Seidman MA, Leeper NJ, Francis GA: Smooth Muscle Cells Contribute the Majority of Foam Cells in Apolipoprotein E-Deficient Mouse Atherosclerosis. Arterioscler Thromb Vasc Biol 2019, 39:876–887.

14. Allahverdian S, Chaabane C, Boukais K, Francis GA, Bochaton-Piallat M-L: Smooth muscle cell fate and plasticity in atherosclerosis. Cardiovasc Res 2018, 114:540–550.

15. Alencar GF, Owsiany KM, Karnewar S, Sukhavasi K, Mocci G, Nguyen AT, Williams CM, Shamsuzzaman S, Mokry M, Henderson CA, Haskins R, Baylis RA, Finn AV, McNamara CA, Zunder ER, Venkata V, Pasterkamp G, Björkegren J, Bekiranov S, Owens GK: Stem Cell Pluripotency Genes Klf4 and Oct4 Regulate Complex SMC Phenotypic Changes Critical in Late-Stage Atherosclerotic Lesion Pathogenesis. Circulation 2020, 142:2045–2059.

16. Cheng P, Wirka RC, Shoa Clarke L, Zhao Q, Kundu R, Nguyen T, Nair S, Sharma D, Kim H-J, Shi H, Assimes T, Brian Kim J, Kundaje A, Quertermous T: ZEB2 Shapes the Epigenetic Landscape of Atherosclerosis. Circulation 2022, 145:469–485.

17. Wirka RC, Wagh D, Paik DT, Pjanic M, Nguyen T, Miller CL, Kundu R, Nagao M, Coller J, Koyano TK, Fong R, Woo YJ, Liu B, Montgomery SB, Wu JC, Zhu K, Chang R, Alamprese M, Tallquist MD, Kim JB, Quertermous T: Atheroprotective roles of smooth muscle cell phenotypic modulation and the TCF21 disease gene as revealed by single-cell analysis. Nat Med, Nature Publishing Group, 2019, 25:1280–1289.

18. Pan H, Xue C, Auerbach BJ, Fan J, Bashore AC, Cui J, Yang DY, Trignano SB, Liu W, Shi J, Ihuegbu CO, Bush EC, Worley J, Vlahos L, Laise P, Solomon RA, Connolly ES, Califano A, Sims PA, Zhang H, Li M, Reilly MP: Single-Cell Genomics Reveals a Novel Cell State During Smooth Muscle Cell Phenotypic Switching and Potential Therapeutic Targets for Atherosclerosis in Mouse and Human. Circulation 2020, 142:2060–2075.

19. Cheng P, Wirka RC, Kim JB, Kim H-J, Nguyen T, Kundu R, Zhao Q, Sharma D, Pedroza A, Nagao M, Iyer D, Fischbein MP, Quertermous T: Smad3 regulates smooth muscle cell fate and mediates adverse remodeling and calcification of the atherosclerotic plaque. Nat Cardiovasc Res 2022, 1:322–333.

20. Owsiany KM, Deaton RA, Soohoo KG, Tram Nguyen A, Owens GK: Dichotomous Roles of Smooth Muscle Cell-Derived MCP1 (Monocyte Chemoattractant Protein 1) in Development of Atherosclerosis. Arterioscler Thromb Vasc Biol 2022, 42:942–956.

21. Gomez D, Baylis RA, Durgin BG, Newman AAC, Alencar GF, Mahan S, St Hilaire C, Müller W, Waisman A, Francis SE, Pinteaux E, Randolph GJ, Gram H, Owens GK: Interleukin-1β has atheroprotective effects in advanced atherosclerotic lesions of mice. Nat Med 2018, 24:1418–1429.

22. Newman AA, Serbulea V, Baylis RA, Shankman LS, Bradley X, Alencar GF, Owsiany K, Deaton RA, Karnewar S, Shamsuzzaman S, Salamon A, Reddy MS, Guo L, Finn A, Virmani R, Cherepanova OA, Owens GK: Multiple cell types contribute to the atherosclerotic lesion fibrous cap by PDGFRβ and bioenergetic mechanisms. Nat Metab 2021, 3:166–181.

23. Shankman LS, Gomez D, Cherepanova OA, Salmon M, Alencar GF, Haskins RM, Swiatlowska P, Newman AAC, Greene ES, Straub AC, Isakson B, Randolph GJ, Owens GK: KLF4-dependent phenotypic modulation of smooth muscle cells has a key role in atherosclerotic plaque pathogenesis. Nat Med 2015, 21:628–637.

24. Nagao M, Lyu Q, Zhao Q, Wirka RC, Bagga J, Nguyen T, Cheng P, Kim JB, Pjanic M, Miano JM, Quertermous T: Coronary Disease-Associated Gene TCF21 Inhibits Smooth Muscle Cell Differentiation by Blocking the Myocardin-Serum Response Factor Pathway. Circ Res 2020, 126:517–529.

25. Cherepanova OA, Gomez D, Shankman LS, Swiatlowska P, Williams J, Sarmento OF, Alencar GF, Hess DL, Bevard MH, Greene ES, Murgai M, Turner SD, Geng Y-J, Bekiranov S, Connelly JJ, Tomilin A, Owens GK: Activation of the pluripotency factor OCT4 in smooth muscle cells is atheroprotective. Nat Med, Nature Publishing Group, 2016, 22:657–665.

26. Kim H-J, Cheng P, Travisano S, Weldy C, Monteiro JP, Kundu R, Nguyen T, Sharma D, Shi H, Lin Y, Liu B, Haldar S, Jackson S, Quertermous T: Molecular mechanisms of coronary artery disease risk at the PDGFD locus. Nat Commun, Nature Publishing Group, 2023, 14:847.

27. Miller CL, Pjanic M, Wang T, Nguyen T, Cohain A, Lee JD, Perisic L, Hedin U, Kundu RK, Majmudar D, Kim JB, Wang O, Betsholtz C, Ruusalepp A, Franzén O, Assimes TL, Montgomery SB, Schadt EE, Björkegren JLM, Quertermous T: Integrative functional genomics identifies regulatory mechanisms at coronary artery disease loci. Nat Commun 2016, 7:12092.

28. Wirka RC, Wagh D, Paik DT, Pjanic M, Nguyen T, Miller CL, Kundu R, Nagao M, Coller J, Koyano TK, Fong R, Woo YJ, Liu B, Montgomery SB, Wu JC, Zhu K, Chang R, Alamprese M, Tallquist MD, Kim JB, Quertermous T: Atheroprotective roles of smooth muscle cell phenotypic modulation and the TCF21 disease gene as revealed by single-cell analysis. Nat Med, Nature Publishing Group, 2019, 25:1280–1289.

29. Kattoor AJ, Pothineni NVK, Palagiri D, Mehta JL: Oxidative Stress in Atherosclerosis. Curr Atheroscler Rep 2017, 19:42.

30. Madamanchi NR, Vendrov A, Runge MS: Oxidative Stress and Vascular Disease. Arteriosclerosis, Thrombosis, and Vascular Biology, American Heart Association, 2005, 25:29–38.

31. Stocker R, Keaney JF: Role of oxidative modifications in atherosclerosis. Physiol Rev 2004, 84:1381–1478.

32. Clempus RE, Griendling KK: Reactive oxygen species signaling in vascular smooth muscle cells. Cardiovasc Res 2006, 71:216–225.

33. Touyz RM, Schiffrin EL: Reactive oxygen species in vascular biology: implications in hypertension. Histochem Cell Biol 2004, 122:339–352.

34. Griendling KK, Sorescu D, Ushio-Fukai M: NAD(P)H Oxidase. Circulation Research, American Heart Association, 2000, 86:494–501.

35. Touyz RM, Schiffrin EL: Reactive oxygen species in vascular biology: implications in hypertension. Histochem Cell Biol 2004, 122:339–352.

36. Vásquez-Vivar J, Duquaine D, Whitsett J, Kalyanaraman B, Rajagopalan S: Altered tetrahydrobiopterin metabolism in atherosclerosis: implications for use of oxidized tetrahydrobiopterin analogues and thiol antioxidants. Arterioscler Thromb Vasc Biol 2002, 22:1655–1661.

37. Xu S, Chamseddine AH, Carrell S, Miller FJ: Nox4 NADPH oxidase contributes to smooth muscle cell phenotypes associated with unstable atherosclerotic plaques. Redox Biology 2014, 2:642–650.

38. Lyle AN, Remus EW, Fan AE, Lassègue B, Walter GA, Kiyosue A, Griendling KK, Taylor WR: Hydrogen Peroxide Regulates Osteopontin Expression through Activation of Transcriptional and Translational Pathways. J Biol Chem 2014, 289:275–285.

39. Griendling KK, Ushio-Fukai M: Redox control of vascular smooth muscle proliferation. The Journal of Laboratory and Clinical Medicine, Elsevier, 1998, 132:9–15.

40. Sundaresan M, Yu ZX, Ferrans VJ, Irani K, Finkel T: Requirement for generation of H2O2 for platelet-derived growth factor signal transduction. Science, New York, N.Y., 1995, 270:296–299.

41. Doanes AM, Irani K, Goldschmidt-Clermont PJ, Finkel T: A requirement for rac1 in the PDGF-stimulated migration of fibroblasts and vascular smooth cells. Biochem Mol Biol Int 1998, 45:279–287.

42. Kang DH, Lee DJ, Kim J, Lee JY, Kim H-W, Kwon K, Taylor WR, Jo H, Kang SW: Vascular Injury Involves the Overoxidation of Peroxiredoxin Type II and Is Recovered by the Peroxiredoxin Activity Mimetic That Induces Reendothelialization. Circulation, American Heart Association, 2013, 128:834–844.

43. Bräutigam L, Jensen LDE, Poschmann G, Nyström S, Bannenberg S, Dreij K, Lepka K, Prozorovski T, Montano SJ, Aktas O, Uhlén P, Stühler K, Cao Y, Holmgren A, Berndt C: Glutaredoxin regulates vascular development by reversible glutathionylation of sirtuin 1. Proc Natl Acad Sci U S A 2013, 110:20057–20062.

44. Choi MH, Lee IK, Kim GW, Kim BU, Han Y-H, Yu D-Y, Park HS, Kim KY, Lee JS, Choi C, Bae YS, Lee BI, Rhee SG, Kang SW: Regulation of PDGF signalling and vascular remodelling by peroxiredoxin II. Nature, Nature Publishing Group, 2005, 435:347–353.

45. Wu Y, Liu Y, Shan L, Li X, Wu B, Guo L: HIF1α/PRDX1 axis drives pulmonary vascular remodeling through DRP1 DeSUMOylation and mitochondrial fragmentation. Biochimica et Biophysica Acta (BBA) - Molecular Basis of Disease 2026, 1872:168094.

46. Park J-G, Yoo J-Y, Jeong S-J, Choi J-H, Lee M-R, Lee M-N, Hwa Lee J, Kim HC, Jo H, Yu D-Y, Kang SW, Rhee SG, Lee M-H, Oh GT: Peroxiredoxin 2 deficiency exacerbates atherosclerosis in apolipoprotein E-deficient mice. Circ Res 2011, 109:739–749.

47. Jeong S-J, Cho MJ, Ko NY, Kim S, Jung I-H, Min J-K, Lee SH, Park J-G, Oh GT: Deficiency of peroxiredoxin 2 exacerbates angiotensin II-induced abdominal aortic aneurysm. Exp Mol Med 2020, 52:1587–1601.

48. Mathai C, Jourd’heuil F, Pham LGC, Gilliard K, Howard D, Balnis J, Jaitovich A, Chittur SV, Rilley M, Peredo-Wende R, Ammoura I, Shin SJ, Barroso M, Barra J, Shishkova E, Coon JJ, Lopez-Soler RI, Jourd’heuil D: Regulation of DNA damage and transcriptional output in the vasculature through a cytoglobin-HMGB2 axis. Redox Biol 2023, 65:102838.

49. Jourd’heuil FL, Xu H, Reilly T, McKellar K, El Alaoui C, Steppich J, Liu YF, Zhao W, Ginnan R, Conti D, Lopez-Soler R, Asif A, Keller RK, Schwarz JJ, Thanh Thuy LT, Kawada N, Long X, Singer HA, Jourd’heuil D: The Hemoglobin Homolog Cytoglobin in Smooth Muscle Inhibits Apoptosis and Regulates Vascular Remodeling. Arterioscler Thromb Vasc Biol 2017, 37:1944–1955.

50. Halligan KE, Jourd’heuil FL, Jourd’heuil D: Cytoglobin is expressed in the vasculature and regulates cell respiration and proliferation via nitric oxide dioxygenation. J Biol Chem 2009, 284:8539–8547.

51. Zweier JL, Ilangovan G: Regulation of Nitric Oxide Metabolism and Vascular Tone by Cytoglobin. Antioxid Redox Signal 2020, 32:1172–1187.

52. Zweier JL, Hemann C, Kundu T, Ewees MG, Khaleel SA, Samouilov A, Ilangovan G, El-Mahdy MA: Cytoglobin has potent superoxide dismutase function. Proc Natl Acad Sci U S A 2021, 118:e2105053118.

53. Zhang S, Li X, Jourd’heuil FL, Qu S, Devejian N, Bennett E, Jourd’heuil D, Cai C: Cytoglobin Promotes Cardiac Progenitor Cell Survival against Oxidative Stress via the Upregulation of the NFκB/iNOS Signal Pathway and Nitric Oxide Production. Sci Rep, Nature Publishing Group, 2017, 7:10754.

54. Pham LGC, Gilliard K, Jourd’heuil F, Mistretta S, Schwarz JJ, Singer HA, Jourd’heuil D: Genetic deletion of cytoglobin exacerbates cardiac hypertrophy and inhibits cardiac fibroblast activation independent of changes in blood pressure. Am J Physiol Heart Circ Physiol 2026, .

55. Mathai C, Jourd’heuil FL, Lopez-Soler RI, Jourd’heuil D: Emerging perspectives on cytoglobin, beyond NO dioxygenase and peroxidase. Redox Biol 2020, 32:101468.

56. Straub AC, Lohman AW, Billaud M, Johnstone SR, Dwyer ST, Lee MY, Bortz PS, Best AK, Columbus L, Gaston B, Isakson BE: Endothelial cell expression of haemoglobin α regulates nitric oxide signalling. Nature, Nature Publishing Group, 2012, 491:473–477.

57. Liu X, El-Mahdy MA, Boslett J, Varadharaj S, Hemann C, Abdelghany TM, Ismail RS, Little SC, Zhou D, Thuy LTT, Kawada N, Zweier JL: Cytoglobin regulates blood pressure and vascular tone through nitric oxide metabolism in the vascular wall. Nat Commun 2017, 8:14807.

58. Hao Y, Stuart T, Kowalski MH, Choudhary S, Hoffman P, Hartman A, Srivastava A, Molla G, Madad S, Fernandez-Granda C, Satija R: Dictionary learning for integrative, multimodal and scalable single-cell analysis. Nat Biotechnol 2024, 42:293–304.

59. Alsaigh T, Evans D, Frankel D, Torkamani A: Decoding the transcriptome of calcified atherosclerotic plaque at single-cell resolution. Commun Biol 2022, 5:1084.

60. Moon KR, Van Dijk D, Wang Z, Gigante S, Burkhardt DB, Chen WS, Yim K, Elzen AVD, Hirn MJ, Coifman RR, Ivanova NB, Wolf G, Krishnaswamy S: Visualizing structure and transitions in high-dimensional biological data. Nat Biotechnol 2019, 37:1482–1492.

61. Van Dijk D, Sharma R, Nainys J, Yim K, Kathail P, Carr AJ, Burdziak C, Moon KR, Chaffer CL, Pattabiraman D, Bierie B, Mazutis L, Wolf G, Krishnaswamy S, Pe’er D: Recovering Gene Interactions from Single-Cell Data Using Data Diffusion. Cell 2018, 174:716–729.e27.

62. Wickham H: ggplot2 [Internet]. Cham, Springer International Publishing, 2016 [cited 2026 Mar 23], . Available from: http://link.springer.com/10.1007/978-3-319-24277-4

63. Yu G, Wang L-G, Han Y, He Q-Y: clusterProfiler: an R Package for Comparing Biological Themes Among Gene Clusters. OMICS: A Journal of Integrative Biology 2012, 16:284–287.

64. Carlson M: org.Mm.eg.db: Genome wide annotation for Mouse. 2024, .

65. Carlson M: org.Hs.eg.db: Genome wide annotation for Human. 2024, .

66. Grootaert MOJ, Bennett MR: Vascular smooth muscle cells in atherosclerosis: time for a re-assessment. Cardiovasc Res 2021, 117:2326–2339.

67. Chappell J, Harman JL, Narasimhan VM, Yu H, Foote K, Simons BD, Bennett MR, Jørgensen HF: Extensive Proliferation of a Subset of Differentiated, yet Plastic, Medial Vascular Smooth Muscle Cells Contributes to Neointimal Formation in Mouse Injury and Atherosclerosis Models. Circ Res 2016, 119:1313–1323.

68. Kumar A, Lindner V: Remodeling with neointima formation in the mouse carotid artery after cessation of blood flow. Arterioscler Thromb Vasc Biol 1997, 17:2238–2244.

69. Aikawa M, Sakomura Y, Ueda M, Kimura K, Manabe I, Ishiwata S, Komiyama N, Yamaguchi H, Yazaki Y, Nagai R: Redifferentiation of smooth muscle cells after coronary angioplasty determined via myosin heavy chain expression. Circulation 1997, 96:82–90.

70. Aikawa M, Sivam PN, Kuro-o M, Kimura K, Nakahara K, Takewaki S, Ueda M, Yamaguchi H, Yazaki Y, Periasamy M: Human smooth muscle myosin heavy chain isoforms as molecular markers for vascular development and atherosclerosis. Circ Res 1993, 73:1000–1012.

71. Gordon D, Reidy MA, Benditt EP, Schwartz SM: Cell proliferation in human coronary arteries. Proc Natl Acad Sci U S A 1990, 87:4600–4604.

72. Herring BP, Hoggatt AM, Griffith SL, McClintick JN, Gallagher PJ: Inflammation and vascular smooth muscle cell dedifferentiation following carotid artery ligation. Physiol Genomics 2017, 49:115–126.

73. Zhang QJ, Goddard M, Shanahan C, Shapiro L, Bennett M: Differential gene expression in vascular smooth muscle cells in primary atherosclerosis and in stent stenosis in humans. Arterioscler Thromb Vasc Biol 2002, 22:2030–2036.

74. Déglise S, Bechelli C, Allagnat F: Vascular smooth muscle cells in intimal hyperplasia, an update. Front Physiol 2023, 13:1081881.

75. Amrute JM, Jung I-H, Yamawaki T, Lin W-L, Bredemeyer A, Diekmann J, Hayat S, Zhang X, Wakefield DL, Luo X, Maryam S, Heo GS, Yang S, Lee CJM, Wang C, Chou C, Kuppe C, Cook KD, Kovacs A, Chintalgattu V, Pruitt D, Barreda J, Stitziel NO, Cheng P, Liu Y, Kramann R, Kreisel D, Foo RS-Y, Rulifson IC, Martin S, Grunert D, Thomas M, Cui J, Quertermous T, Bengel FM, Jackson S, Li C-M, Ason B, Lavine KJ: Targeting modulated vascular smooth muscle cells in atherosclerosis via FAP-directed immunotherapy. Science, New York, N.Y., 2026, :eadx1736.

76. Chen R, McVey DG, Shen D, Huang X, Ye S: Phenotypic Switching of Vascular Smooth Muscle Cells in Atherosclerosis. J Am Heart Assoc 2023, 12:e031121.

77. Ou L, Li X, Chen B, Ge Z, Zhang J, Zhang Y, Cai G, Li Z, Wang P, Dong W: Recombinant Human Cytoglobin Prevents Atherosclerosis by Regulating Lipid Metabolism and Oxidative Stress. J Cardiovasc Pharmacol Ther 2018, 23:162–173.

