## Supplementary Material for "Smooth Muscle Cell Cytoglobin is a Negative Regulator of Atherosclerotic Fibrous Cap Development"

**Supplemental Figures**

### Supplemental Figure 1

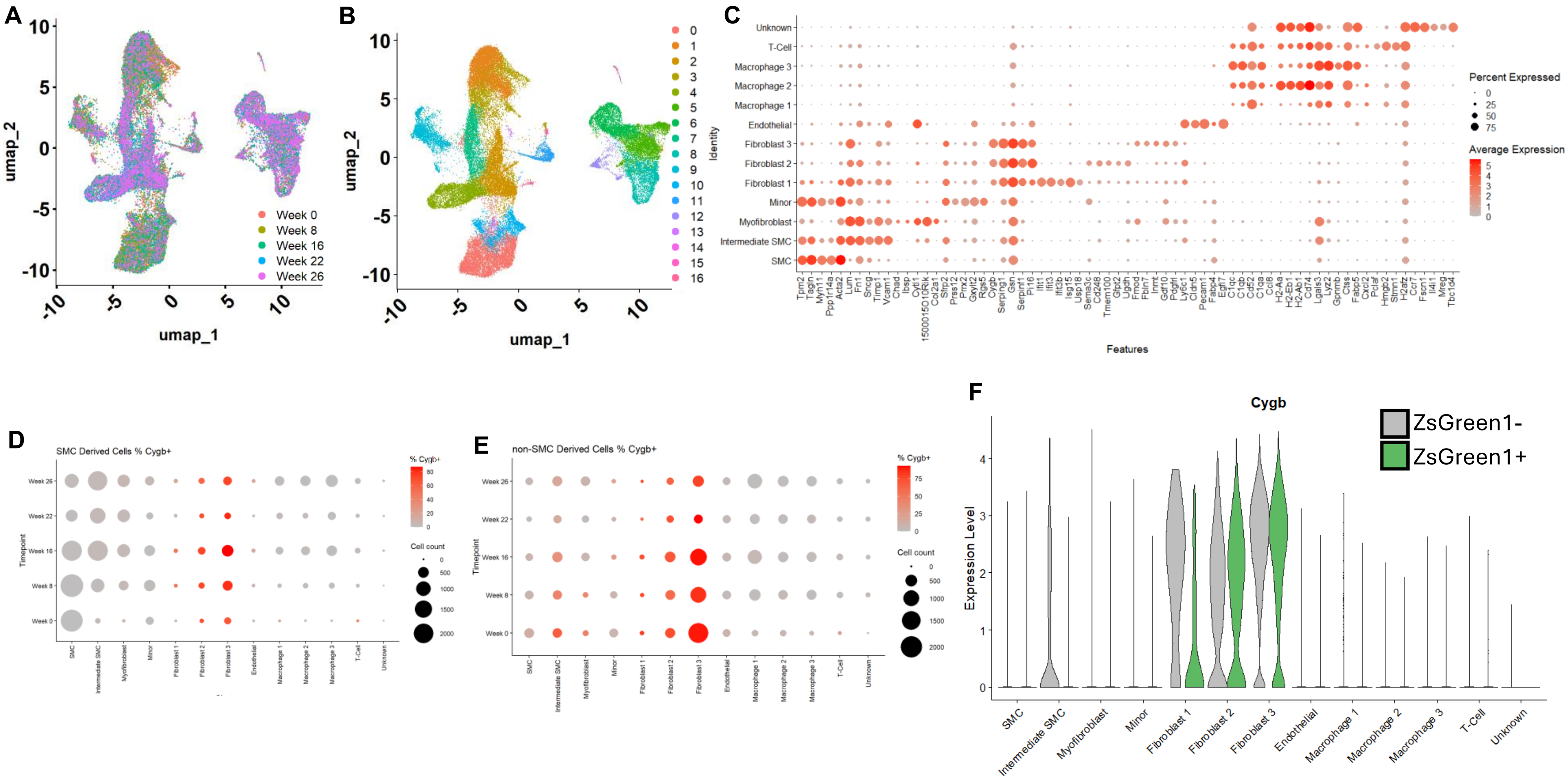


**Supplemental Figure 1: A)** Integrated UMAP projections identifying timepoints of western diet feeding from publicly available single cell dataset (GSE155514). **B)** UMAP visualizing 17 transcriptionally distinct cell clusters. **C)** Dot plot of traditional markers for cell identity. **D-E)** Dot plot showing CYGB expression by Western diet timeline in lineage traced ZsGreen1+ cells **(D)** and ZsGreen1- cells **(E)**. Dot size represents cells count, color represents percentage of cells expressing Cygb. **F)** Violin plots representing expression of Cygb between cell types in ZsGreen1+ and ZsGreen1- cells.

#### Supplemental Figure 2

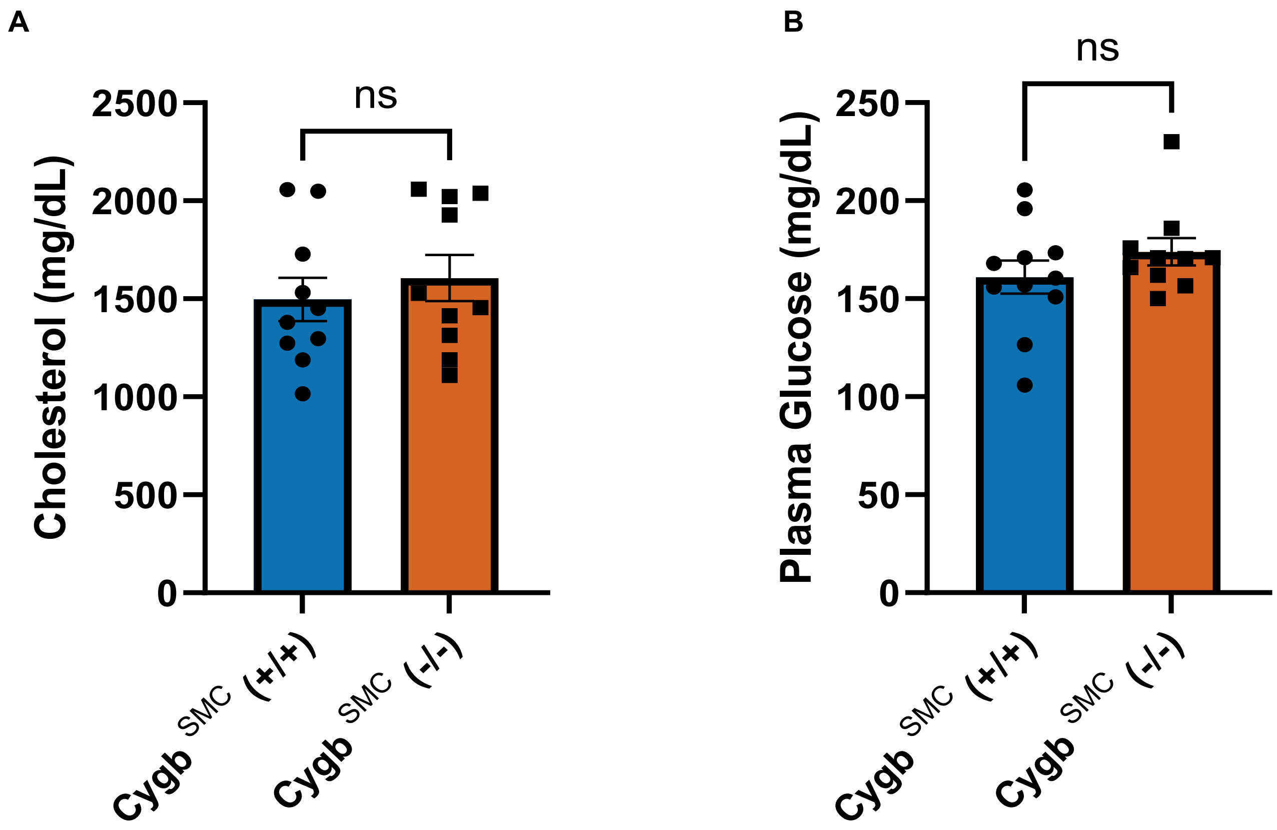


**Supplemental Figure 2: A)** Bar graphs showing mouse plasma cholesterol levels measured six hours after fasting and three weeks after PCSK9-AAV8-GOF virus and western diet (mg/dl). **B)** Bar graph showing plasma cholesterol 16 weeks after PCSK9-AAV8-GOF virus and western diet (mg/dL). Results are represented as mean ± SEM and each dot represent an individual mouse. Student's t-test was used. ns p> .05, * p< 0.05, ** p< 0.01, *** p< 0.001.

### Supplemental Figure 3

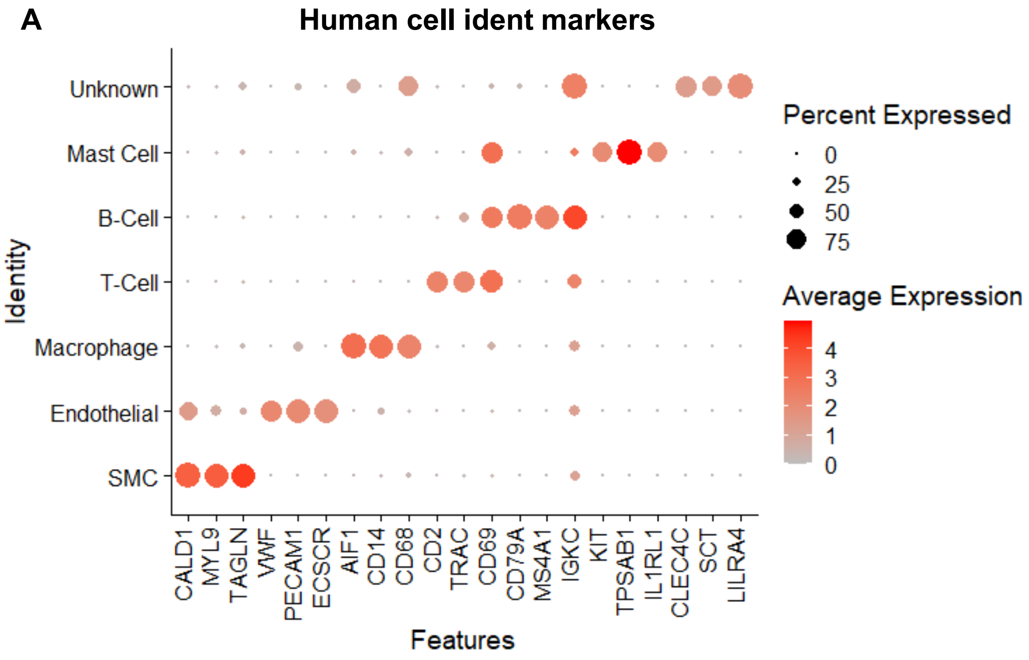


**Supplemental Figure 3: A)** Dot plot showing markers for identify cell identity.

### Supplemental Table 1

| Plaque Stability | Calcification | Race | Sex | Age |
|---|---|---|---|---|
| Stable | No | W | M | 47 |
| Very Stable | No | W | F | 37 |
| Stable | Yes | W | M | 59 |
| Very Stable | No | B | M | 54 |
| Very Stable | No | B | M | 32 |
| Unstable | No | W | M | 50 |
| Unstable | No | W | M | 57 |
| Unstable | Yes | B | F | 52 |
| Unstable | No | B | M | 39 |
| Unstable | Yes | W | M | 59 |
| No Plaque | No | W | M | 50 |
| No Plaque | No | W | F | 38 |
| No Plaque | No | B | M | 40 |
| No Plaque | No | B | M | 37 |
| No Plaque | No | W | M | 49 |

**Supplemental Table 1:** Clinical and histopathological characteristics of human atherosclerotic plaque samples used for staining analyses.

#### **Supplemental Table 2.** Supplies and reagents

| Reagents | | |
|---|---|---|
| Acetic Acid | Sigma | 320099 |
| Agarose | Thermo Scientific | R0492 |
| Bluing Reagent (Epredia, Richard Allan Scientific 7301) | Fisher | 22-050-116 |
| Boving Serum Albuman | Sigma | A2153 |
| Citrate Buffer | Sigma | C9999-1000 |
| Clarifying reagent (Epredia, Richard Allan Scientific 7401) | Fisher | 22-050-116 |
| DAPI | Sigma | D9542-5mg |
| DirectPCR (Tail) | Viagen | 102-T |
| DNA Ladder -Trackit 100 bp | Invitrogen | 10488058 |
| Eosin Y (Epredia, Richard Allan Scientific 7111) | Fisher Scientific | 22-050-110 |
| Ethanol (Ethyl alcohol - pure) | Sigma | E7023 |
| Formaldehyde solution 37% | Sigma | 252549 |
| Glycine | Sigma | G8898 |
| Goat Serum | Vector labs | S-1000 |
| GoTaq G2 master mix | Promega | M7823 |
| Hematoxylin (Epredia, Signature Series 7211) | Fisher Scientific | 22-050-111 |
| 2-Methyl Butane | Sigma | M32631 |
| Mounting Media (Epredia Signature Series 4112) | Fisher Scientific | 22-110-610 |
| OCT compound | Tissue-Tek | 4583 |
| mPCSK9 AAV8-D377Y | Vector Biolabs | SKU VB-377Y |
| Phosphate buffered Saline (DPBS) | Corning | 21-031-CV |
| Proteinase K | Viagen | 501-PK |
| Saline | Baxter | 2F7123 |
| Sucrose | Sigma | S1888 |
| Sunflower Seed Oil | Sigma | S5007 |
| SYBR safe DNA gel stain | Invitrogen | S33102 |
| Tamoxifin | Sigma | T5648 |
| Triology | Sigma | 920-P06 |
| Triton X-100 | Sigma | T9284 |
| Vectashield Vibrance mounting media | Vector labs | H-1700 |
| Western Diet | Inotiv | TD88137 |
| Xylene | Sigma | 534056 |

| Kits/Materials | | |
|---|---|---|
| Cholesterol Kit | FUJIFILM Healthcare Americas Corp | 999-02601 |
| Colorfrost plus microscope slides | Cardinal health | M6148-3P |
| Cryomolds | Electron Microscopy Sciences | 62534-10 |
| Masson's Trichrome Stain Kit | Polyscience, Inc | 25088-1 |
| Glucose detection strips | Accu-Chek Avida Plus | 6908268001 |
| Glucose monitor | Accu-Chek Avida | 8340331001 |

| Antibodies | | |
|---|---|---|
| Alpha smooth muscle actin (ACTA2) | Invitrogen | MA5-11547 |
| Cytoglobin Rabbit (CYGB) | Sigma | HPA017757 |
| Goat anti Rabbit Alexafluor 594 | Invitrogen | A32740 |
| Goat anti Rabbit Alexafluor 647 | Invitrogen | A21236 |
| Mouse IgG isotype control | Vector labs | I-2000 |
| Rabbit IgG isotype Control | Vector labs | I-1000 |
